# SETD7-mediated lysine monomethylation is abundant on non-hyperphosphorylated nuclear Tau

**DOI:** 10.1101/2020.01.14.905786

**Authors:** Maria Bichmann, Nuria Prat Oriol, Ebru Ercan-Herbst, David C. Schöndorf, Borja Gomez Ramos, Vera Schwärzler, Per Haberkant, Laura Gasparini, Dagmar E. Ehrnhoefer

## Abstract

Human tauopathies including Alzheimer’s disease (AD) are characterized by alterations in the post-translational modification (PTM) pattern of Tau, leading to the formation of insoluble aggregates, neuronal dysfunction and degeneration. Using a mass spectrometry approach, we identified multiple sites of lysine monomethylation on Tau isolated from a detergent-soluble fraction of human brain, some of which were increased in early AD samples. Brain tissues derived from a mouse model of tauopathy demonstrate an age-dependent increase in methylation at specific sites, with methylated Tau enriched in the soluble nuclear fraction and not associated with hyperphosphorylated, insoluble Tau species. Furthermore, we show that the protein lysine methyltransferase SETD7 methylates Tau at K132 and demonstrate an interaction with K130, an additional methylation site in close vicinity. These findings shed light on the function of a novel type of PTM on Tau that provide a potential signal for its translocation to different subcellular sites. Since the mislocalization and depletion of Tau from axons is associated with tauopathies, our findings may furthermore provide insight into this disease-associated phenomenon.

## Introduction

Tau is a neuronal protein that is subject to multiple types of post-translational modifications (PTMs), among which phosphorylation is the most studied. Tau PTMs are regulated developmentally (Yu, Run et al., 2009) and are altered in a number of neurodegenerative diseases, the so-called tauopathies (Sergeant, Delacourte et al., 2005). Familial mutations in Tau are associated only with a small subset of tauopathies, thus, the underlying cause for Tau dysfunction is unclear for most sporadic diseases including Alzheimer’s disease (AD). In AD, Tau accumulates intraneuronally in detergent-insoluble aggregates with fibrillar morphology, termed neurofibrillary tangles (NFT). Tau in these NFTs is heavily modified, predominantly by hyperphosphorylation and truncation events (Quinn, Corbett et al., 2018, Wang & Mandelkow, 2016). NFTs are considered to be end-stage products of a prolonged process of Tau misfolding, oligomerization and aggregation and soluble misfolded Tau species have been proposed to cause significant neuronal dysfunction before the appearance of mature NFTs (Choi & Gandhi, 2018, Lasagna-Reeves, Castillo-Carranza et al., 2012).

Tau is a predominantly cytoplasmic protein that localizes to axons in mature neurons, where it stabilizes the microtubule cytoskeleton. However, Tau mislocalizes to soma and dendrites in tauopathy (Ittner & Ittner, 2018). Furthermore, nuclear or perinuclear localization of Tau has been described (Frost, Hemberg et al., 2014, Maina, Bailey et al., 2018, Sotiropoulos, Galas et al., 2017). Tau does not contain a nuclear import or localization signal, but its translocation to non-axonal compartments may be regulated by differential PTM patterns. For example, Tau mis-sorting to the somatodendritic compartment can be modulated by acetylation (Sohn, Tracy et al., 2016). However, no PTM-dependent mechanism has been demonstrated to date for the nuclear translocation of Tau.

To better understand the status of Tau PTMs in health and disease, we have previously performed an ELISA-based analysis of detergent-soluble extracts derived from human brain tissues of controls and early stage Alzheimer’s disease patients (Braak stages 0 - IV) (Ercan-Herbst, Schöndorf et al., 2019). This study was necessarily biased towards commonly studied PTMs, for which well-validated antibodies are available (Ercan, Eid et al., 2017). Using an unbiased mass spectrometry (MS)-based method (Behrendt, Bichmann et al., 2019), we now report novel PTM events on detergent-soluble Tau extracted from human brain tissues, including a set of previously unknown sites for lysine monomethylation, and determine that a subset of these PTMs is increased in early stage AD brains. We also show that monomethylation at specific lysine sites increases with age in a mouse model of tauopathy, and our data suggest that methylation is enriched in a pool of soluble, non-hyperphosphorylated Tau. We further demonstrate that the protein lysine methyltransferase (PKMT) SETD7 regulates Tau methylation at two sites in close vicinity, and these modified Tau species preferentially localize to nuclear fractions *in vitro* and *in vivo*.

## Results

### Monomethylation is abundant on soluble Tau in human brain

Published MS-based studies on the PTM landscape of Tau in AD mostly rely on the purification of insoluble protein fractions, which predominantly contain NFTs, and are often performed on brain samples derived from end-stage AD patients (Braak stages V-VI) (Funk, Thomas et al., 2014, Hanger, Byers et al., 2007, Mair, Muntel et al., 2016). However, evidence suggests that Tau dysfunction is already present at earlier stages and may be mediated by soluble Tau species (Tracy & Gan, 2018). We recently developed a method to enrich detergent-soluble Tau from human brain for MS-based analyses of Tau (Behrendt et al., 2019). This method relies on immunoprecipitation from a Triton X-soluble fraction with three antibodies directed against different regions in the protein and uses two parallel protease digests with either trypsin or AspN to generate complementary Tau peptides, achieving a sequence coverage of 80 – 97% (Fig. 1A). We analyzed entorhinal cortex samples from donors classified as Braak stages 0-I as well as Braak stages III-IV to represent non-affected controls and early (often still clinically asymptomatic) AD, respectively (Table 1). This led to the identification of 12 phosphorylation sites which have been previously described, one novel acetylation site (Table 2) and 10 monomethylation sites, including 6 novel sites (K130, K150, K294, K298, K343, K438) and 4 described previously (K132, K259, K353, K385 (Funk et al., 2014, Huseby, Hoffman et al., 2019)). The majority of these methylated lysines is located in the repeat domains (Fig. 1B), but also a C-terminal site (meK438) and two sites in the projection domain (meK130 and meK132) were consistently modified in multiple brain samples.

**Table 1:** Characteristics of human brain samples used in the study.

| Sample | Age | Sex | Braak stage | PMI [h] | Cause of death | Source |
| --- | --- | --- | --- | --- | --- | --- |
| Control 1 | 77 | M | 0 | 17.45 | Carcinomatosis, cancer of unknown primary | Bristol brain bank |
| Control 2 | 72 | M | I | 16.25 | Multiple myeloma, paroxysmal atrial fibrillation | Bristol brain bank |
| Control 3 | 96 | F | I | 72 | Urinary sepsis, paroxysmal atrial fibrillation, hypertension | King's college London |
| Control 4 | 78 | M | I | 30.5 | Metastatic renal carcinoma, bilateral pulmonary emboli | King's college London |
| Control 5 | 87 | M | I | 48 | Cardiac failure | King's college London |
| Control 6 | 78 | M | 0 | 56 | Metastatic brain tumor | Bristol brain bank |
| Control 7 | 86 | F | I | 38.5 | Cerebrovascular disease | Bristol brain bank |
| Control 8 | 96 | F | I | 49 | Cerebrovascular disease | Bristol brain bank |
| Control 9 | 92 | M | I | 56,5 | Cerebral amyloid angiopathy | Bristol brain bank |
| AD 1 | 80 | M | III | 50 | Bronchopneumonia, idiopathic pulmonary fibrosis | Bristol brain bank |
| AD 2 | 81 | M | IV | 38 | Lewy body dementia | Bristol brain bank |
| AD 3 | 83 | F | III | 70.75 | Neutropenic sepsis, endometrial carcinoma, ischaemic heart disease | Bristol brain bank |
| AD 4 | 95 | F | IV | 87.5 | Frailty of old age | Bristol brain bank |
| AD 5 | 91 | F | III | 27.5 | Frailty of old age, cerebrovascular disease, atrial fibrillation | Bristol brain bank |
| AD 6 | 95 | M | III | 25.45 | Corticobasal degeneration | Bristol brain bank |
| AD 7 | 85 | F | III | 13.5 | Old age | Bristol brain bank |
| AD 8 | 95 | F | III | 65.5 | Alzheimer's disease | Bristol brain bank |
| AD 9 | 82 | M | III | 51.25 | Alzheimer's disease | Bristol brain bank |

**Table 2:**
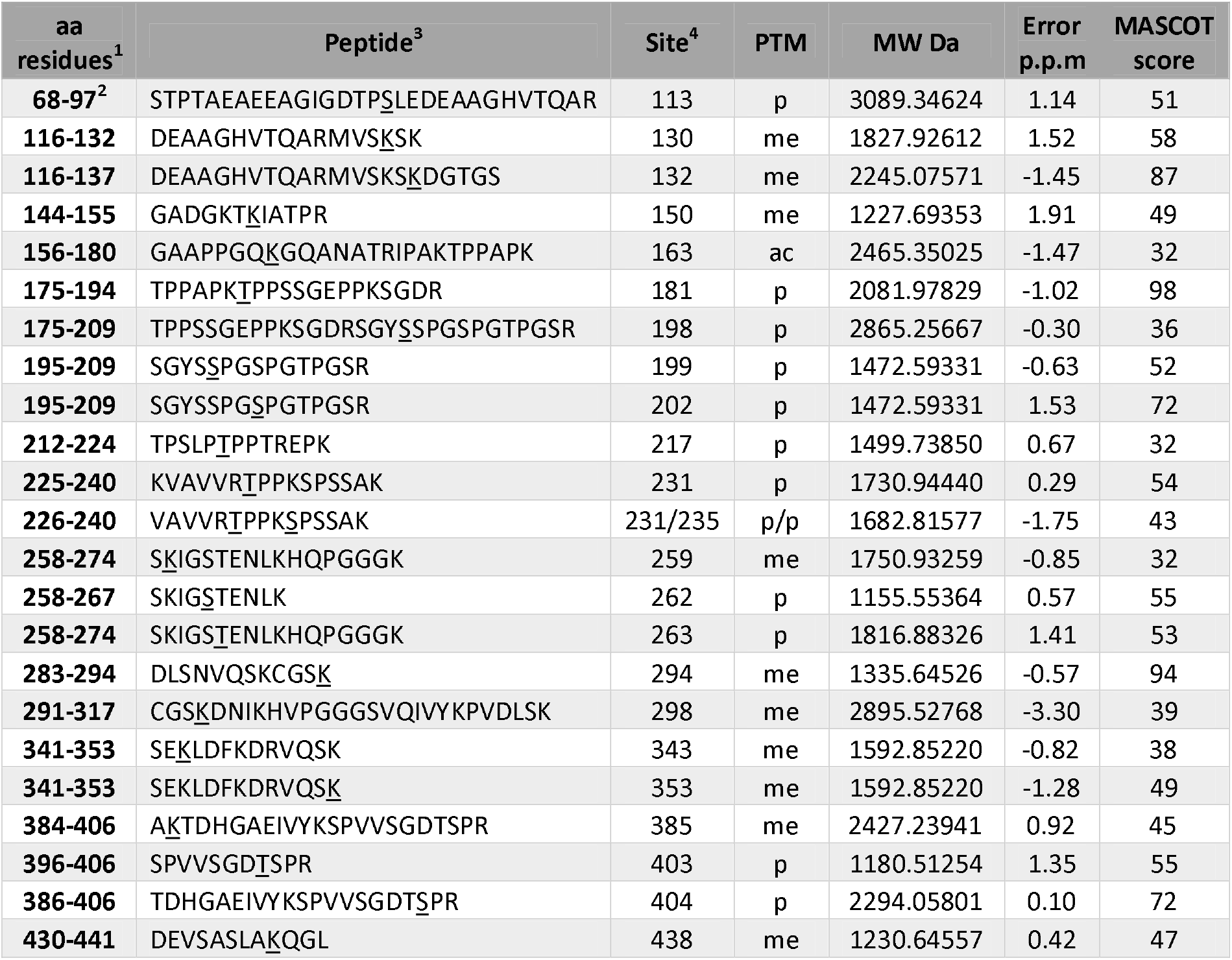
PTM modifications identified on tau protein immunoprecipitated from human entorhinal cortex. Trypsin and AspN digests of tau were analyzed by LC-MS/MS and resulting m/z data was analyzed by MASCOT using 10 ppm mass tolerance for the search. ^1^amino acid (aa) residue numbering conforms to human 2N4R tau, except for peptide 68-97 (pS113). ^2^aa residue according to human 1N4R tau. ^3^<u>K</u> modified lysine, <u>S</u> modified serine, <u>T</u> modified threonine. modified site according to human 2N4R tau. me: monomethylation, ac: acetylation, p: phosphorylation, MW: molecular weight in Dalton.

**Fig. 1:**
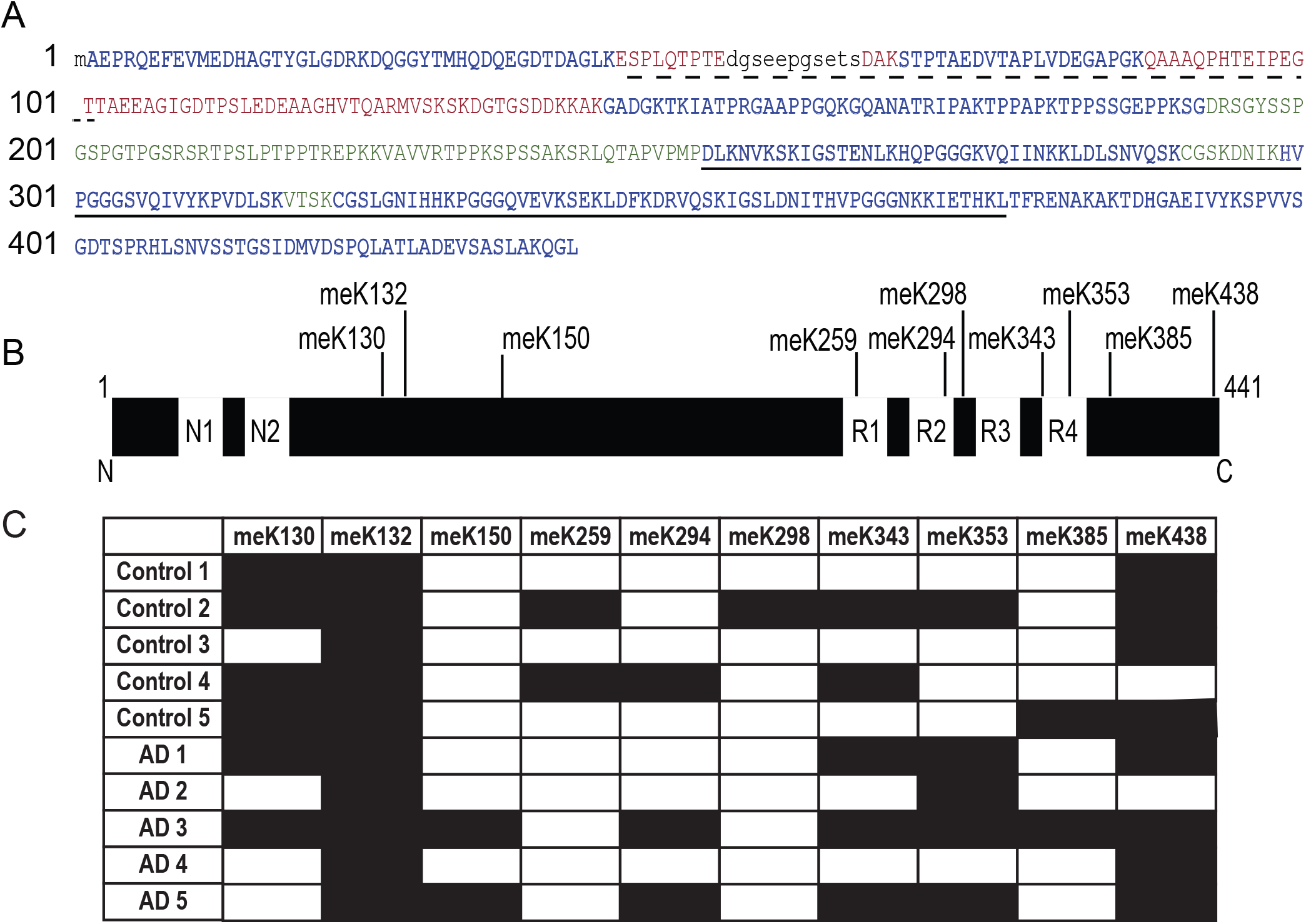
Multiple sites of lysine monomethylation detected on soluble Tau from human entorhinal cortex tissue. **(A)** Sequence coverage of the human 2N4R Tau isoform (UniProtKB-P10636-8). Dashed line: N-terminal repeats N1 and N2, solid line: microtubule binding domain (MTBR). Font color in blue: sequence covered by peptides obtained by both AspN and trypsin digestion, red: sequence covered exclusively by AspN-derived peptides, green: sequence covered exclusively by trypsin-derived peptides, black lower case: sequence not recovered by MS. **(B)** Summary of lysine monomethylation sites identified by LC-MS/MS on 2N4R Tau. **(C)** Prevalence of each monomethyl-modified lysine across all human brain samples analyzed.

### Generation and validation of methyl-Tau specific antibodies

To analyze differences in PTM abundance between control and AD brains, we developed an ELISA-based method using methyl-Tau specific antibodies. First, we raised antibodies against the five most prevalent monomethyl sites (identified in a total of 5 or more individual brain samples): meK130, meK132, meK343, meK353 and meK438 (Fig. 1B and C).

In order to verify that the novel antibodies are specific for methylated Tau, we performed dot blot assays using recombinant Tau peptides spanning the different modification sites with or without a mono-methyl modification. In addition, we used recombinant full-length Tau protein (2N4R isoform), which was subjected to reductive methylation, a process that chemically modifies all available lysine residues with methyl groups (Funk et al., 2014). All antibodies were highly specific for the methylated peptide against which they were raised and recognized chemically methylated full-length Tau (Suppl. Fig. S1). To test for site specificity in the context of intracellularly expressed Tau, we used transfections in HEK cells of either wild-type (wt) Tau (2N4R) or mutants at the different methyl sites (Suppl. Fig. S2). Western blotting confirmed the specificity of meK130, meK132 and meK353 Tau antibodies. Both the meK343 and the meK438 Tau antibodies also detected the mutant protein in this experiment and thus appeared to be non-specific with respect to their methylation site (Suppl. Fig. S2). We therefore focused on meK130, meK132 and meK353 for all antibody-based downstream analyses.

### Tau methylation is altered in human AD and in a tauopathy mouse model

A recent study has suggested that Tau lysine methylation is altered in the human brain with age and differences potentially exist between control and AD patients (Huseby et al., 2019). To assess whether our sites of interest are differentially methylated between Braak 0-I and Braak III-IV patient brain samples, we used a sensitive electrochemiluminescence ELISA method using commercially available total Tau antibody-coated plates for capture and our novel methyl antibodies as detection antibodies. We found that the abundance of both meK130 and meK353 modifications were increased in the soluble fraction of Braak III-IV entorhinal cortices, while the meK132 antibody yielded a similar signal for both sample groups (Fig. 2A).

**Fig. 2:**
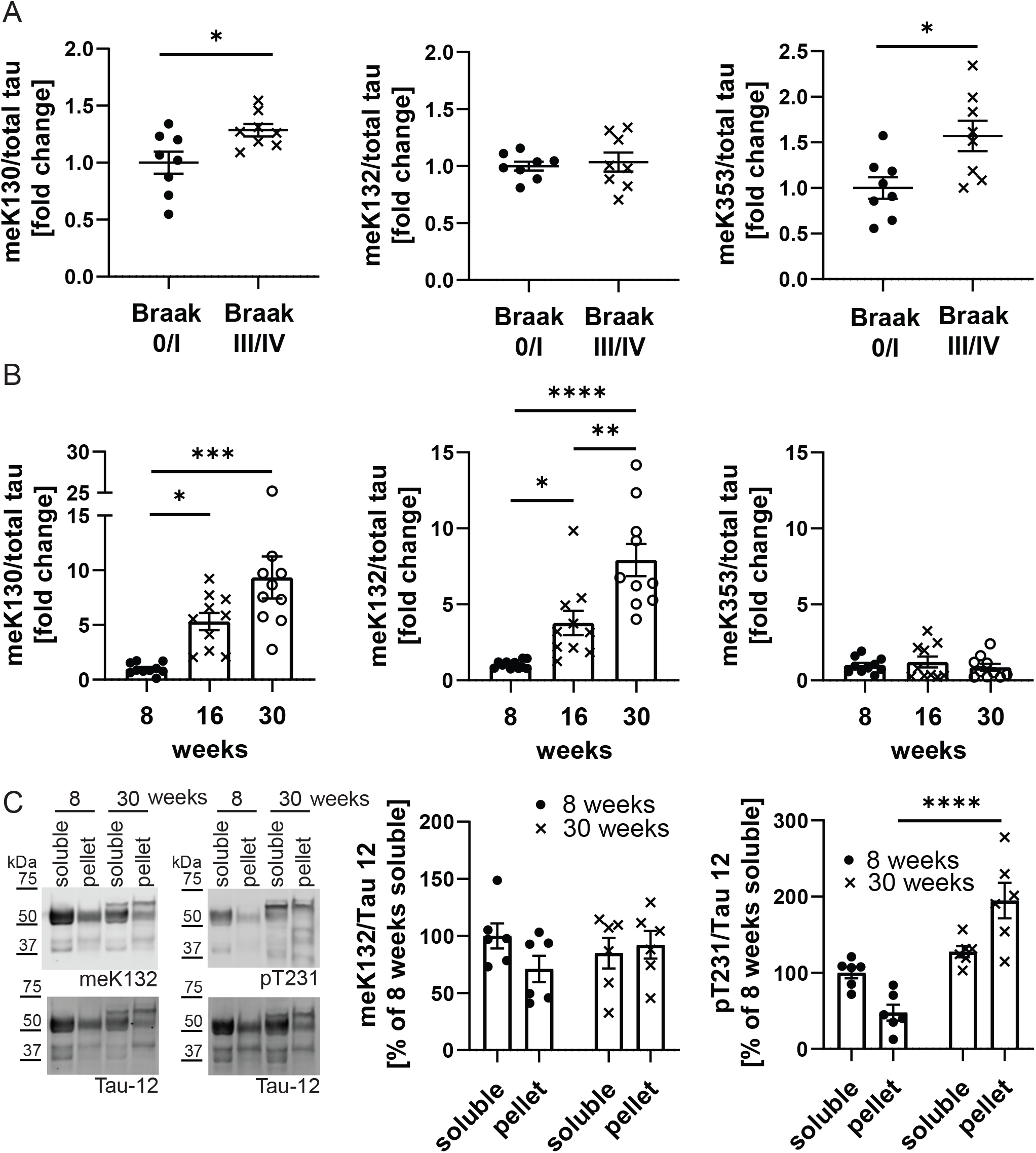
Tau methylation is increased in human AD patient brains and increases with age in a mouse model of tauopathy. **(A)** Electrochemiluminescence ELISA demonstrates an increase of meK130 and meK353-modified Tau in human entorhinal cortex samples at Braak stage III/IV. Data are normalized to total Tau levels obtained by BT2 ELISA. Statistical significance was determined by Mann Whitney test, *: p <0.05. **(B)** Electrochemiluminescence ELISA demonstrates an age-dependent increase in meK130 and meK132-modified Tau in tg4510 mouse hippocampus. Data are normalized to total Tau levels obtained by Tau 5 ELISA. Statistical significance was determined by one-way ANOVA (meK130: ANOVA p=0.0002, meK132: ANOVA p<0.0001), Tukey’s multiple comparisons test was used for post-hoc analysis. *: p<0.05, **: p<0.01, ***: p< 0.001, ****: p< 0.0001. **(C)** Triton-X soluble and insoluble fractions from tg4510 mouse hippocampi were analyzed at different ages by Western blotting. While pT231-modified Tau accumulates with age in the pellet fraction, the solubility of meK132-modified Tau does not change with age. Statistical significance was determined by two-way ANOVA (meK132: age p=0.8029, solubility p=0.3776; pT231: age p<0.0001, solubility p=0.5981), Tukey’s multiple comparisons test was used for post-hoc analysis. ****: p< 0.0001.

Next, we investigated whether these changes are recapitulated in a transgenic mouse model of tauopathy, the rTg4510 mice (Ramsden, Kotilinek et al., 2005). rTg4510 mice progressively accumulate sarkosyl-insoluble Tau and exhibit intraneuronal Tau inclusions throughout the forebrain from 4 months of age (Santacruz, Lewis et al., 2005). In the Triton-soluble hippocampal fraction from mice aged 2-6 months, all three Tau methylation sites were detected by ELISA, but only meK130 and meK132 signals progressively increased with age (Fig. 2B).

At the 30 weeks timepoint, rTg4510 mice exhibit abundant Tau insoluble inclusions (Santacruz et al., 2005). We therefore asked whether we might underestimate the amount of methylated Tau by focusing on the soluble fraction. We solubilized the pellets obtained after Triton-extraction by boiling in SDS and compared the relative abundance of methylated and phosphorylated Tau in both soluble and insoluble (pellet) fractions. While phosphorylation at T231 (pT231) accumulated in the pellet fraction in 6-month old rTg4510 mice, both meK130 and meK132 did not (Fig. 2C). Taken together, these data suggest that although methylated Tau accumulates with age in rTg4510 mouse brain samples, it does not preferentially localize to insoluble, aggregated Tau species that are hyperphosphorylated and accumulate in parallel.

### Methylated Tau preferentially localizes to the nucleus

Monomethylation is a small, charge-neutral PTM that can mediate nuclear translocation of the target protein, which is a common mechanism to regulate the activity of transcription modulators (Cornett, Ferry et al., 2019). We therefore decided to investigate the subcellular localization of meK130- and meK132-Tau. We found that the majority of Tau protein in iPSC-derived neurons localized to the cytoplasmic and the soluble nuclear fractions and was not apparently bound to DNA (Fig. 3A). Interestingly, both meK130- and meK132-Tau were strongly enriched in the soluble nuclear fraction in these samples (Fig. 3B), suggesting that methylated Tau is preferentially located to the neuronal nucleus.

**Fig. 3:**
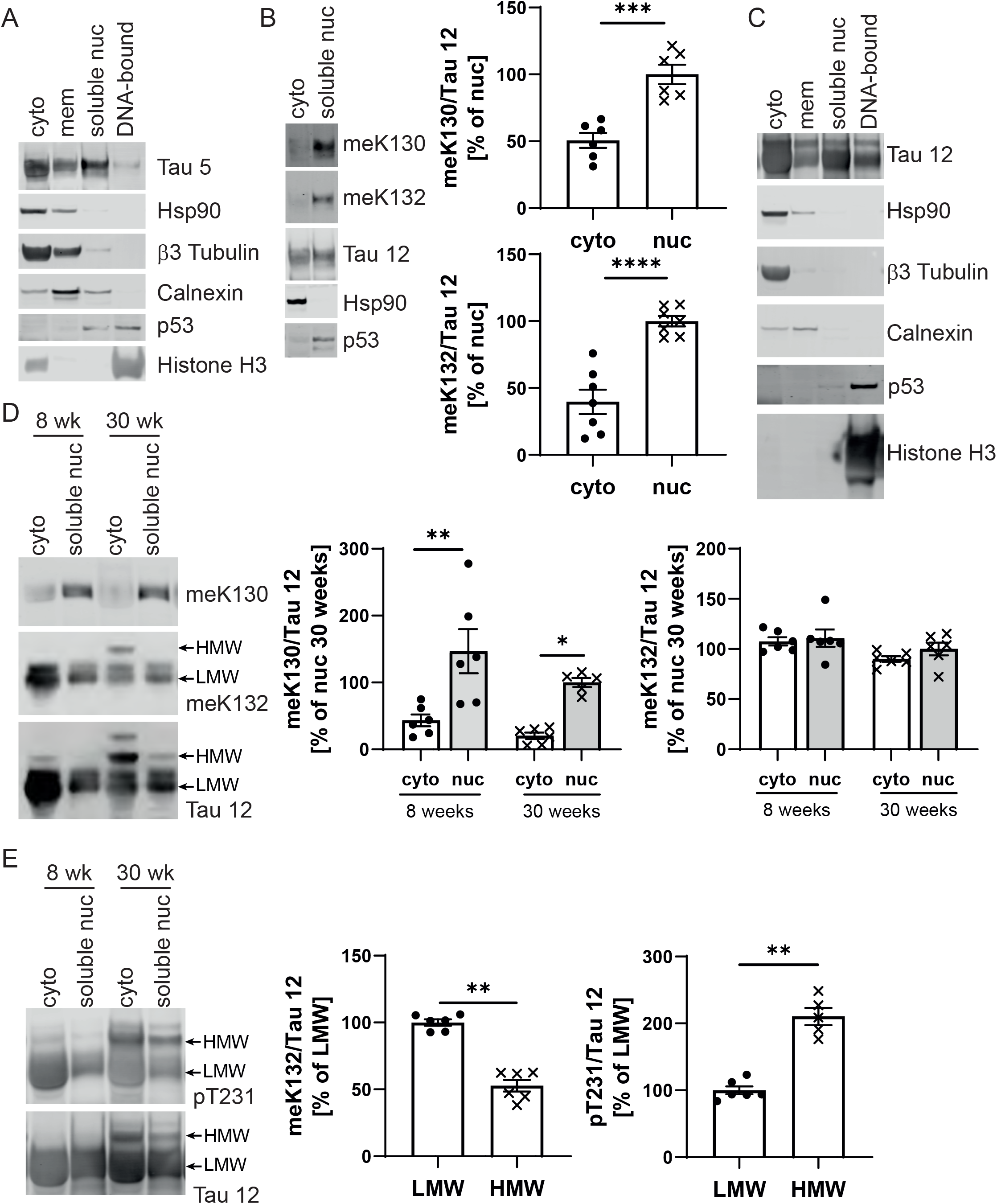
Methylated Tau preferentially localizes to the nucleus. **(A)** iPSC-derived neurons were subjected to subcellular fractionation and the distribution of Tau was analyzed by Western blotting. The purity of fractions was determined by analyzing the distribution of marker proteins. As demonstrated by a representative blot, the soluble nuclear fraction contains abundant Tau protein but is free from the cytosolic protein Hsp90 and the Tau interactor β3 tubulin. **(B)** Representative Western blots and quantification demonstrate that meK130 and meK132-modified Tau preferentially localize to the soluble nuclear fraction in iPS-derived neurons. **(C)** Cortical tissue from 30 weeks old tg4510 mice was subjected to subcellular fractionation and the distribution of Tau was analyzed by Western blotting. The purity of fractions was determined by analyzing the distribution of marker proteins. Similar to in vitro data, the soluble nuclear fraction contains abundant Tau protein but is free from the cytosolic protein Hsp90 and the Tau interactor β3 tubulin. **(D)** Representative Western blots and quantification demonstrate that meK130, but not meK132-modified Tau preferentially localizes to the soluble nuclear fraction in both young and old tg4510 cortex. **(E)** pT231-modified Tau, but not meK132-modified Tau is enriched in the hyperphosphorylated HMW Tau band. Cytosolic fractions were quantified, representative Western blot and quantification are shown. Statistical significance in **(B)** and **(E)** was determined by Mann Whitney test, **: p<0.01, ***: p<0.001, ****: p<0.0001. Statistical significance in **(D)** was determined by two-way ANOVA (meK130: age p=0.0721, fraction p<0.0001; meK132: age p=0.0264, fraction p=0.2756), Tukey’s multiple comparisons test was used for post-hoc analysis. *: p<0.05, **: p< 0.01.

We next analyzed the subcellular distribution of Tau in cortical brain extracts from rTg4510 mice. Similar to cultured neurons, total Tau was enriched in both cytoplasmic and the soluble nuclear fractions (Fig. 3C). In these samples, meK130-Tau was exclusively present in the soluble nuclear fraction, while Tau methylated at K132 was more evenly distributed between fractions with only a non-significant trend towards increased nuclear localization at 6 months of age (Fig. 3D). Additionally, extracts from 6-month old mice exhibited high molecular weight (HMW) Tau bands, which stained strongly for a phospho-epitope present in hyperphosphorylated Tau (pT231, Suppl. Fig. S3). This band was abolished upon treatment with λ-phosphatase, confirming that the shift in molecular weight is caused by hyperphosphorylation (Suppl. Fig. S3). Comparing the abundance of meK130 and meK132 antibody immunoreactivity for low-molecular weight (LMW) and HMW Tau, we found that HMW Tau is significantly less methylated than the LMW species (Fig. 3E). This effect was particularly striking for meK130, which was not detected in the HMW band. In contrast, the analysis of pT231 modified Tau showed an enrichment in the HMW band, as expected. Together with our previous observation that only phosphorylated, but not methylated Tau accumulates in the insoluble fraction (Fig. 2C), this suggests that methylation is independent from hyperphosphorylation events.

### Inhibition or knockdown of SETD7 reduces Tau methylation at meK130 and meK132

While a few published studies have reported different lysine methylation events on Tau, the enzymes involved in these processes are unknown (Funk et al., 2014, Huseby et al., 2019, Thomas et al., 2012). We analyzed the Tau sequence for potential recognition sites for protein lysine methyltransferases and identified the consensus motif for SETD7 (SET Domain Containing 7) at aa130-132 (R/KSK(me), (Dhayalan, Kudithipudi et al., 2011)). SETD7 is a lysine methyltransferase with several cytoplasmic and nuclear non-histone substrates, which have implicated the enzyme in a variety of conditions including cancer and inflammation (Batista & Helguero, 2018). Taking advantage of a selective inhibitor of SETD7, (R)-PFI-2 (Barsyte-Lovejoy, Li et al., 2014), we investigated whether SETD7 induces Tau methylation. Treating SH-SY5Y neuroblastoma cells stably expressing GFP-0N4R Tau with (R)-PFI-2 significantly reduced both meK132-Tau and meK130-Tau, while the amount of meK353-Tau remained unaltered (Fig. 4A-B). The treatment did not reduce total Tau levels (Suppl. Fig. S4A and B). Consistently, we also observed a significant reduction of endogenous meK132- and meK130-Tau in iPS-derived neurons after treatment with (R)-PFI-2 (Fig. 4C-D). Again, total Tau levels remained unaffected by the treatment (Suppl. Fig. S4C and D).

**Fig. 4:**
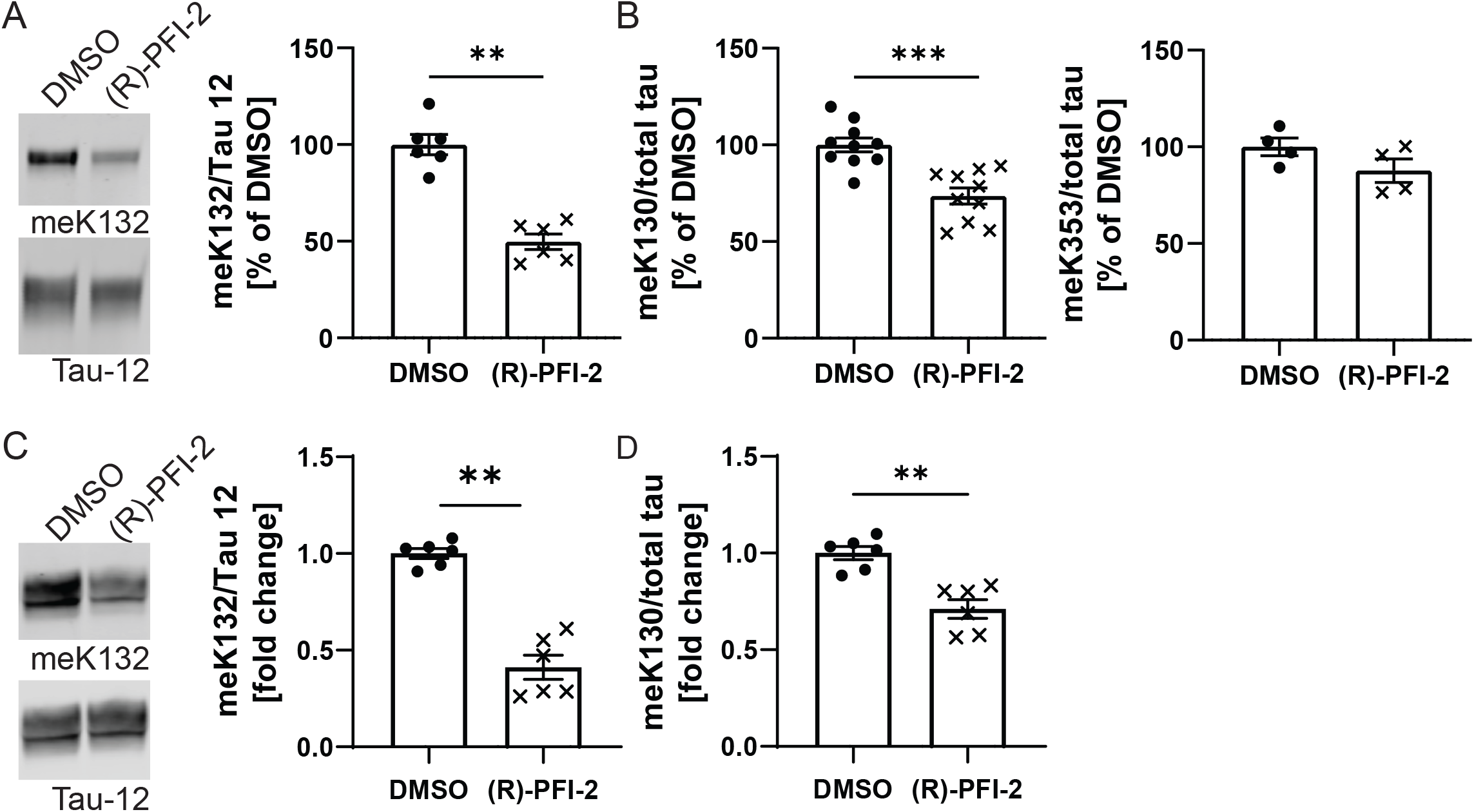
The SETD7 inhibitor (R)-PFI-2 reduces Tau methylation at K130 and K132. **(A)** Representative Western blots and quantification demonstrate a significant decrease in meK132-modified Tau after treatment of SHSY5Y cells with 5 μM (R)-PFI-2 for 24h. **(B)** Electrochemiluminescence ELISA shows that treatment with 5 μM (R)-PFI-2 for 24h reduces the levels of meK130, but not meK353 Tau in SH-SY5Y cells. **(C)** Western blots and quantification demonstrate a significant decrease in meK132-modified Tau after treatment of iPS-derived neurons with 5 μM (R)-PFI-2 for 48h. **(D)** Electrochemiluminescence ELISA shows that treatment with 5 μM (R)-PFI-2 for 48h reduces the levels of meK130 Tau in iPS-derived neurons. Statistical significance for all panels was determined by Mann Whitney test. **: p<0.01, ***: p< 0.001.

To exclude potential off-target effects of the pharmacological inhibitor, we decreased SETD7 levels by shRNA-mediated knockdown in SH-SY5Y cells. Using a mix of two lentiviral shRNA constructs, we obtained a reduction of SETD7 protein of app. 90% (Fig. 5A). This was accompanied by reduced levels of both meK132- and meK130-Tau in these cells (Fig. 5B and C), confirming the importance of SETD7 for Tau methylation at these residues. Unlike treatment with the pharmacological inhibitor, shRNA mediated knockdown of SETD7 also led to a reduction in total Tau levels, possibly due to the longer time-course of the shRNA experiment and the strong reduction of SETD7 protein (Suppl. Fig. S4E).

**Fig. 5:**
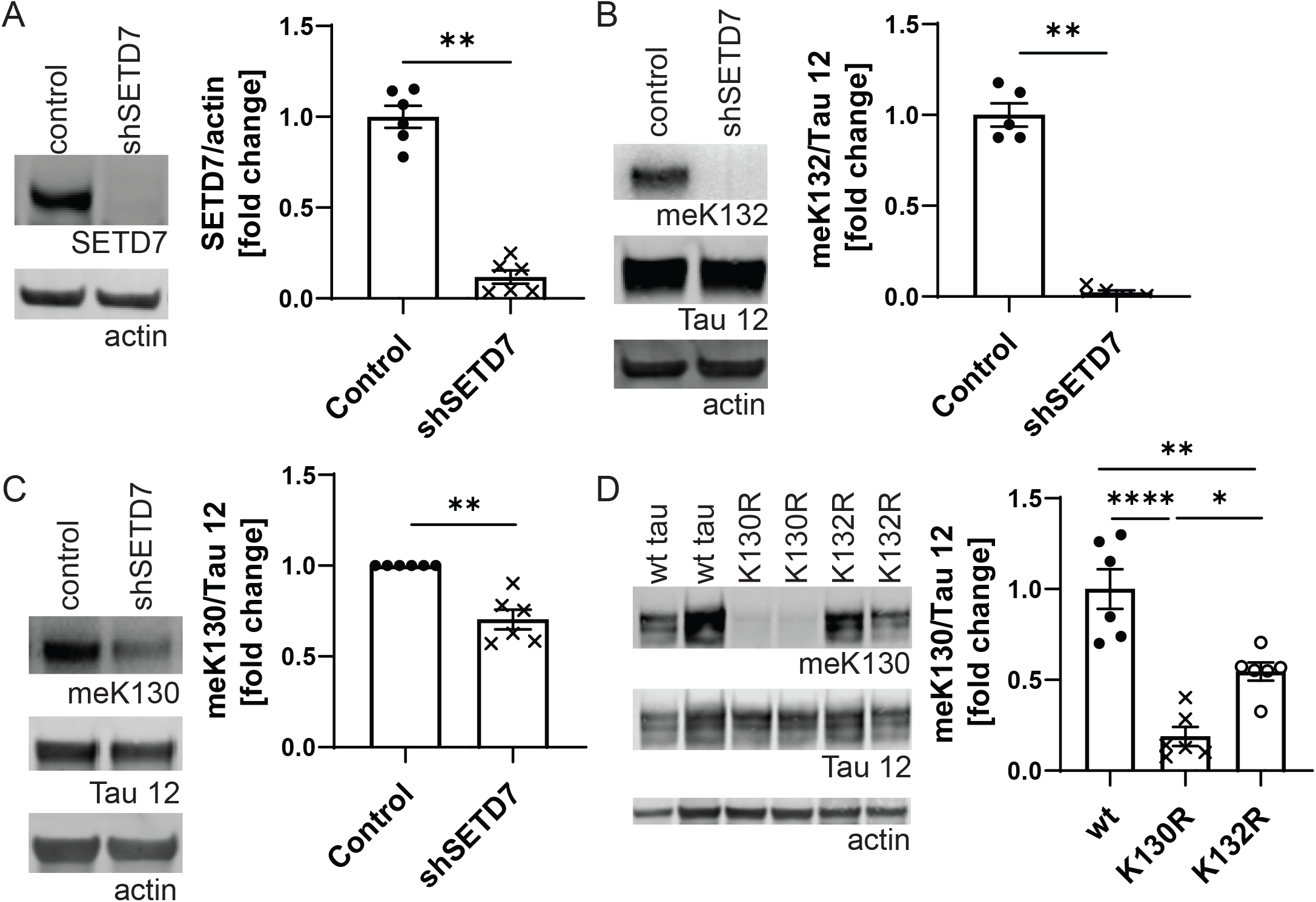
shRNA-mediated knockdown of SETD7 expression reduces meK130 and mek132-modified Tau in SHSY5Y cells. **(A)** Representative Western blots and quantification demonstrate a strong reduction of SETD7 protein in SHSY5Y cells treated with shRNA. **(B)** meK132-modified Tau is absent in SH-SY5Y cells treated with shRNA against SETD7, representative Western blots and quantification. **(C)** Representative Western blots and quantification demonstrate that shRNA against SETD7 also reduces the levels of meK130-modified Tau in SH-SY5Y cells, albeit to a smaller degree than the meK132 modification. **(D)** Transient transfection of wt, K130R and K132R mutant Tau 2N4R into HEK cells demonstrates that the K132R mutation significantly decreases meK130-modified Tau levels. Representative Western blots and quantification are shown. Statistical significance in **(A)**, **(B)** and **(C)** was determined by Mann Whitney test, **: p<0.01. Statistical significance in **(D)** was determined by one-way ANOVA (ANOVA p<0.0001), Tukey’s multiple comparisons test was used for post-hoc analysis. *: p<0.05, **: p<0.01, ****: p< 0.0001.

For both the inhibitor and the shRNA treatment, the reduction in methylation was always more pronounced for K132 than K130. The K130 site does not exactly match the consensus sequence for SETD7-mediated methylation (VSK(me) instead of R/KSK(me), (Dhayalan et al., 2011)). It is therefore possible that K130 is not directly targeted by SETD7, or that methylation at this site requires the initial modification of K132 by SETD7. To investigate the mutual interaction of these sites, we assessed meK130 in the presence of a K132R mutation, which abolishes methylation at K132 (Fig. 5D and suppl. Fig. S4F). In agreement with our hypothesis we found that the levels of the meK130 modification were significantly reduced for K132R Tau, suggesting that methylation at these sites may indeed be hierarchically regulated and that the inhibition of SETD7 may indirectly reduce the levels of meK130.

## Discussion

While recent mass spectrometry studies have described lysine methylation on the Tau protein in human brain samples derived from both AD patients and non-affected controls (Funk et al., 2014, Huseby et al., 2019, Thomas et al., 2012), protein lysine methyltransferases involved in this process or the functional consequences of Tau methylation in health and disease were so far unexplored. Here we show that Tau methylation at selected sites is more abundant in human AD brain samples compared to controls and increases with age in the rTg4510 mouse model of tauopathy. The specific sites that exhibit differential methylation only partially overlap between human and mouse brains. Tau methylation at K130 is increased in both species, while meK132-Tau is not different between human Braak 0-I and Braak III-IV brain tissue samples but increases with age in rTg4510 mice. meK353 on the other hand is altered in human AD, but not in rTg4510 brain. These differences may be due to species-dependent regulation of enzymes involved in Tau methylation, or the overlay of methylation patterns for endogenous mouse and transgenic human Tau in the mouse model. Furthermore, aging-related changes may be observed in the longitudinally sampled mice but not in human samples, which were all obtained from elderly donors. K353 in particular is a residue that can be subject to additional PTMs, namely dimethylation, ubiquitination or acetylation, which may be regulated differently in the two species (Cook, Carlomagno et al., 2014, Cook, Gendron et al., 2012, Cripps, Thomas et al., 2006, Frost et al., 2014, Thomas et al., 2012).

The Tau sequence preceding the methylation site at K132 corresponds to the consensus sequence for the protein lysine methyltransferase (PKMT) SETD7 (Dhayalan et al., 2011). Using an inhibitor as well as an shRNA-mediated knockdown approach we confirmed that this enzyme is responsible for the generation of the meK132 modification. Interestingly, inhibition or knockdown of SETD7 also reduced Tau methylation at K130, even though this site does not correspond to the SETD7 consensus sequence (VSK(me) instead of R/KSK(me)). The reduction of meK130 was also consistently less pronounced compared to the effect of SETD7 inhibition on meK132, suggesting that meK130 may not be a direct target. We also show that the K132R mutation, which abolishes methylation at K132, also reduces the meK130 signal, supporting the hypothesis that the methylation events at these two sites are linked. This observation, together with our mass spectrometry data and published findings demonstrating several additional lysine methylation sites in Tau, suggest that the Tau methylome forms a complex regulatory system similar to the regulation of p53 through differential methylation at adjacent sites by multiple PKMTs (Wu, Connolly et al., 2017).

Tau is subject to a large number of other PTMs, most notably phosphorylation (Wang & Mandelkow, 2016). Hyperphosphorylation is strongly associated with the formation of aggregated and insoluble forms of Tau that accumulate in AD patients over the course of the disease, a process that is recapitulated in the rTg4510 mouse model (Ramsden et al., 2005). Interestingly, we found that lysine methylation is less abundant on high-molecular weight, hyperphosphorylated Tau and does not accumulate in the insoluble pellet fraction of 6-month old rTg4510 mice. These findings are in agreement with a published in vitro study suggesting that lysine methylation renders Tau resistant to the formation of fibrils observed by electron microscopy (Funk et al., 2014). Methylated Tau also showed a different subcellular distribution pattern compared to phosphorylated Tau, with a preferential localization to the soluble nuclear fraction in biochemical studies. In particular meK130-Tau was more abundant in the nuclear fraction in both iPS-derived neurons and mouse brain samples, suggesting that this PTM may be important for targeting Tau to the nucleus.

Promoting nuclear localization of target proteins is a known function of lysine methylation, and has been reported among others also for non-histone targets of SETD7 such as p53 or the estrogen receptor (Chuikov, Kurash et al., 2004, Subramanian, Jia et al., 2008). Most lysine methylation sites so far identified on Tau are situated in the microtubule binding (MTB) domain (Funk et al., 2014, Huseby et al., 2019), and methylation may thus contribute to lower affinity of Tau for microtubules and facilitate its re-distribution to other subcellular compartments. Interestingly, a recent study reports preferential nuclear localization of an N279K Tau mutant associated with frontotemporal dementia, for which the novel lysine also falls within the MTB region (Ritter, Avila et al., 2018). Further studies are however warranted to determine the overall association of Tau lysine methylation at multiple sites with its subcellular localization and the potential impact of altered methylation patterns in human tauopathies.

## Materials and Methods

### Human brain samples

Anonymized human post-mortem tissue was obtained from the London Neurodegenerative Diseases Brain Bank and the Southwest Dementia Brain Bank, members of the Brains for Dementia Research Network. Donor characteristics are described in Table 1.

### Transgenic mouse brain samples

Heterozygous rTg4510 mice (licensed from the Mayo Clinic, Jacksonville Florida, USA) (Santacruz et al., 2005) were used in the study. All transgenic mice were bred for AbbVie by Charles River Laboratories (Sulzfeld, Germany). The mice were in a temperature- and humidity-controlled room with a 12:12 hour dark/light cycle with ad *libitum* access to water and food. All animal experiments were performed in full compliance with the Principles of Laboratory Animal Care (NIH publication No. 86-23, revised 1985) in an AAALAC accredited program where veterinary care and oversight was provided to ensure appropriate animal care. All animal studies were approved by the government of Rhineland Palatinate (Landesuntersuchungsamt Koblenz) and conducted in accordance with the directive 2010/63/EU of the European Parliament and of the Council on the protection of animals used for scientific purpose, the ordinance on the protection of animals used for experimental or scientific purposes (German implementation of EU directive 2010/63; BGBl. I S. 3125, 3126), the Commission Recommendation 2007/526/EC on guidelines for the accommodation and care of animals used for experimental and other scientific purposes, the German Animal Welfare Act (BGBl. I S. 1206, 1313) amended by Art. 1 G from 17 December 2018 I 2586. Tg4510 mice express 0N4R human P301L mutant hTau under the CaMKIIα promoter. For this study, mice were culled and hippocampi and cortices were harvested from 2-, 4- and 6-month old heterozygous rTg4510 mice and stored at −80°C until use.

### Brain lysate preparation

Both mouse and human samples were homogenized in Triton lysis buffer (150 mM NaCl, 20 mM Tris pH7.5, 1 mM EDTA, 1 mM EGTA, 1 % Triton-X100 and protease, phosphatase, demethylase (500 μM IOX1, 2 μM Daminozide, 10 μM Paragyline Hydrochloride), deacetylase (10 μM Trichostatin A, 5 mM Nicotinamide), O-GlcNAcase (1 μM Thiamet-G) inhibitors) with a Dounce homogenizer (Carl Roth). The tubes were twice scraped over a hard surface with in-between incubation on ice for 10 min. Samples were centrifuged for 20 min at 16 000x g at 4°C, and the protein concentration in the supernatant was determined with a BCA assay (BioRad, cat. No. 5000112).

For mouse brain, Triton-insoluble pellets were resuspended in sample buffer (65 mM Tris-HCl, 5% Glycerol, 1% SDS) and boiled for 15 min at 95°C, followed by centrifugation for 25 min at 16 000x g, 4°C. The supernatant containing Triton-insoluble Tau was used for subsequent Western blot analyses.

### Immunoprecipitation and sample preparation for LC-MS/MS

250 μg protein from each human brain sample lysate was used for immunoprecipitation (IP) with a combination of Tau12 (Biolegend, cat. no. 806501), Tau 5 (Abcam, cat. no. ab80579) and HT7 (Thermo Fisher, cat. no. MN1000) as described previously (Behrendt et al., 2019). Briefly, immunoprecipitated samples were eluted with 50 mM glycine HCl pH 2.8, separated by SDS-PAGE and Coomassie-stained bands were excised. Samples were subjected to either an in-gel tryptic- or to an in-gel AspN digest, and peptides were separated using nanoAcquity UPLC (Waters) with a nanoAcquity trapping (nanoAcquity Symmetry C18, 5μm, 180 μm × 20 mm) and analytical column (nanoAcquity BEH C18, 1.7μm, 75μm × 200mm), which was coupled to an LTQ Orbitrap Velos Pro (Thermo Fisher) using the Proxeon nanospray source as previously described (Behrendt et al., 2019).

### Methyl- Tau antibodies

Immunization and antibody purification were performed by Innovagen (Lund, Sweden). Antibodies were raised in rabbits against the respective methylated peptides indicated in Suppl. Fig. S1. The animals were subjected to a total of three booster injections, with the final immune serum harvest 2 months after the first immunization. Immune sera were affinity purified against the appropriate methyl-peptide, and the purified antibodies were stored in aliquots in PBS at −80°C until use.

### Recombinant Tau purification

Full length Tau 2N4R was expressed and purified as described previously (Behrendt et al., 2019). Briefly, E. coli BL21(DE3) cells (Sigma, cat. no. CMC0014) transformed with a pET19n vector containing 2N4R Tau were induced by the addition of 1 mM IPTG and harvested by centrifugation. Cleared lysates in 50 mM Na-phosphate pH 7.0, 1 mM EGTA, 1 mM DTT, cOmplete protease inhibitors (Roche), benzonase (Merck) and 10 μg/ml lysozyme (Sigma) were boiled for 20 min at 100°C. Supernatants were loaded onto a combination of a HiTrap Q and a HiTrap SP column (GE Healthcare) and eluted in a gradient to running buffer containing 300 mM NaCl. Tau-containing fractions were further purified with a HiLoad 16/600 Superdex 75 pg size exclusion chromatography column (GE Healthcare). Fractions were analyzed by SDS-PAGE, pooled according to purity, flash-frozen in liquid nitrogen and stored at −80°C.

### In vitro methylation of recombinant Tau

100 μg of recombinant Tau 2N4R were incubated for 2h at room temperature in 100 mM sodium citrate pH 6 (Sigma, cat. no. W302600), 100 mM sodium cyanoborohydride (Sigma, cat. no. 156159) and 20 mM formaldehyde (Fisher Scientific, cat. no. 28908) with a final volume of 100 μl. 10 μl of 550 mM glycine (Roth, cat. no. 3187) were added, and Zeba Spin desalting columns (Thermo Fisher, cat. no. 87767) were used according to manufacturer’s instructions to remove excessive sodium cyanoborohydride.

### Immunoblotting

To generate dot blots for testing the specificity of methyl-Tau antibodies, 3 μg methylated peptides, bare peptides, *in vitro* methylated full-length Tau (2N4R) and recombinant Tau (2N4R) were spotted onto nitrocellulose membranes.

For Western blots, 25 μg total protein were boiled at 95°C for 5 min in sample buffer, separated on a 10 % SDS-PAGE and immunoblotted onto PVDF membranes.

The membranes were blocked with blocking buffer (5% BSA in TBST (Roth)) and incubated with primary antibodies overnight at 4°C: Tau12, mouse, 1:500, (Biolegend, cat. no. 806501); Tau5, mouse, 1:1000, (Abcam, cat. no. ab80579); meK130, rabbit, 1:500, (INNOVAGEN); meK132, rabbit, 1:1000 (INNOVAGEN); meK343, rabbit, 1:1000 (INNOVAGEN); meK353, 1:200, (INNOVAGEN); meK438, rabbit, 1:1000 (INNOVAGEN); Actin, rabbit 1:1000, (Cell Signaling Technology, cat. no. 4910); SETD7, mouse, 1:1000, (Thermo Fisher, cat. no. 730055)). Antibodies were diluted in 5% BSA in 1X TBST (1X TBS, 0.05% Tween-20). The next day, secondary antibody incubations (1:20000, IRDye Donkey anti-mouse 800 and IRDye Donkey anti-rabbit 680) were done in blocking solution for one hour at room temperature. The membranes were then washed three times with 1X TBST, once with 1X TBS and imaged on a Li-Cor Odyssey CLx scanner.

For weak signals, anti-rabbit HRP-linked antibody was used (Cell Signaling Technology, 1:5000) in blocking buffer for one hour at RT. The membranes were then washed three times with 1X TBST, once with 1X TBS. ECL solution clarity Max (Bio-Rad, cat. no. 1705062) was added to the membrane and chemiluminescent signals were detected on a Fusion FX7 (Vilber).

### Electrochemiluminescence ELISA

To quantify methylated Tau in human brain samples, total Tau was captured onto ELISA plates supplied with the V-Plex Human Total Tau Kit (Meso Scale Discovery, Cat. No: K151LAE) according to the manufacturer’s protocol. Briefly, plates were incubated for 1 hr at RT on a plate shaker with 100 μL MSD Eluent per well, washed three times with Tris wash buffer (50 mM Tris-HCl pH 7.5, 150 mM NaCl and 0.02 % Tween-20), 2 μg lysate in 50 μL TBS was added and incubated for 1 hr at RT on a plate shaker. Plates were washed three times with Tris wash buffer and incubated for 1 hr at RT on a plate shaker with 25 μl of 2 μg/ml detection antibody (methyl Tau antibodies (INNOVAGEN) or BT2 (Thermo Fisher, cat. no. MN1010) both labeled with MSD Sulfo-Tag-NHS-Ester (Meso Scale Discovery, cat. no. R31AA) and diluted in 1% Blocker A solution). Plates were washed three times with Tris wash buffer and read with the MESO QuickPlex SQ 120 (Meso Scale Discovery) after a 5 min incubation in 150 μL 1X Read Buffer (Meso Scale Discovery, cat. no. R92TC).

For mouse brain samples and SHSY5Y cells, Gold Streptavidin small-spot 96-well plates (Meso Scale Discovery, cat. no. L45SA) were blocked with 5% (w/v) Blocker A solution in Tris wash buffer for 1 hr at RT on a plate shaker. Plates were washed three times with Tris wash buffer and coated with 1 μg of biotinylated antibody diluted in 25 μL of 1% Blocker A solution for 1 hr at RT on a plate shaker. Antibody biotinylation was performed according to manufacturer’s instructions (EZ-Link Sulfo-NHS-Biotin, Thermo Scientific, cat. no. 21217), after BSA removal with the Melon Gel IgG Purification Kit (Thermo Scientific, cat. no 45212) if necessary. For mouse brain samples biotin Tau5 (Abcam) was used as a total Tau antibody, for SHYSY5Y cells biotin HT7 (Thermo Fisher, cat. no. MN1000) was used. Antibody-coated plates were washed three times with Tris wash buffer, 5 μg of protein lysates (diluted to 50 μl with TBS) were added and incubated for 1 hr at RT on a plate shaker. Plates were washed three times with Tris wash buffer and incubated for 1 hr at RT on a plate shaker with 25 μl of 1 μg/ml detection antibody (for mouse samples: HT7; for SHSY5Y samples: Tau12, both labeled with MSD Sulfo-Tag-NHS-Ester, Meso Scale Discovery) diluted in 1 % Blocker A solution. Plates were washed three times with Tris wash buffer and 150 μl of 2X Read Buffer were added five minutes before the signal was measured on a MESO QuickPlex SQ 120 (Meso Scale Discovery).

### Culture and transfection of cell lines

HEK 293T cells were cultured at 37°C in 5% CO_2_ in DMEM+ GlutaMax (Thermo Fisher, cat. no. 10566016) with 10% fetal bovine serum (Sigma, cat. no. F7524-500ml) and 1% penicillin/streptomycin (Fisher Scientific, cat. no. 15307583). Cells were used for transfections between passage 10-20. Transfection was performed at a cell confluency of 60 – 70% using JetPrime reagent (PolyPlus) according to manufacturer’s instructions. For each construct, 1 μg DNA (wt 2N4R Tau, K130R 2N4R Tau, K132R 2N4R Tau, K343R 2N4R Tau, K353R 2N4R Tau and K438R 2N4R Tau) was used. Tau constructs contained an N-terminal HA tag and were cloned into the pcDNA3 vector, mutants were generated by site-directed mutagenesis (Q5 site directed mutagenesis, New England BioLabs, cat. no. E0552S) and verified by sequencing. Cells were transfected and incubated for 24 h before they were harvested by scraping, centrifuged for 5 min at 2000× g at 4°C, washed once with ice-cold 1X PBS and centrifuged again for 5 min at 2000x g at 4°C. Samples were stored at −20°C until further use.

SHSY5Y neuroblastoma cells stably expressing 0N4R Tau (Innoprot, cat. no. P30722-4R) were cultured at 37°C in 5% CO_2_ in RPMI 1640 + GlutaMax media (Thermo Fisher, cat. no. 61870044) with 10% fetal bovine serum and 250 μg/mL G418 (Fisher Scientific, cat. no. 10131035). Cells were used for experiments between passage 3-20. 5 μM (R)-PFI-2 (Sigma cat. no. SML1408) was added to the cells at a confluency of 50%, followed by an incubation for 16 h. Cells were harvested by scraping, centrifuged for 5 min at 2000× g at 4°C, washed once with ice-cold 1X PBS and centrifuged for 5 min at 2000× g at 4°C. Samples were stored at −20°C until further use.

### Neuron culture

Neurogenin 2 (NGN2) induced neurons were differentiated as previously described with minor modifications (Wang, Ward et al., 2017). A doxycycline inducible NGN2 expression cassette was stable integrated in the AAV1 locus using TALEN technology by Bioneer (Denmark). iPSCs were split in a concentration of 100 000 cells/cm^2^ on matrigel (BD) coated plates in mTesR (Stem Cell Technologies). At day 1 after splitting, the medium was changed to N2/B27 medium (50% DMEM/F12, 50 % Neurobasal, 1:200 N2, 1:100 B27, 1% PenStrep, 0.5 mM Non-essential amino acids, (all Invitrogen), 50 μM ß-mercaptoethanol (Gibco), 2.5 μg/ml insulin and 1 mM sodium pyruvate (both Sigma)) with 2 μg/ml doxycycline (Sigma). The medium was changed daily. On day 4, cells were split with accutase (Invitrogen) and re-seeded in a density of 200 000 cells/cm^2^ in N2/B27 medium with doxycycline and 10 μM Rock inhibitor Y-27632 (Selleckchem) on matrigel coated plates in the final format. N2/B27 with doxycycline was changed daily until day 7. On day 8 the medium was switched to final maturation medium (FMM; N2/B27 with 20 ng/ml BDNF, 10 ng/ml GDNF (both Peprotech), 1 mM dibutyryl-cAMP (Sigma) and 200 μM ascorbic acid (Sigma)). The medium was changed every third day until cells were used for treatment with (R)PFI-2 at day 21. N2/B27 medium with DMSO or 5 μM (R)-PFI-2 (Barsyte-Lovejoy et al., 2014) were added and refreshed once after 24 h. 48 h after initial treatment, cells were detached with ice-cold 1X PBS and centrifuged for 5 min at 2000x g at 4°C. Samples were stored at −20°C until further use.

### SETD 7 Knock down

shRNA sequences against SETD7 were designed using an online tool (www.biosetta.com) and synthesized (Eurofins). Oligonucleotide sequences were as follows:

shRNA-1:

top: 5′ – ACCGGGCCAGGGAGTTTACACTTAGTTAATATTCATAGCTAAGTGTAAACTCCCTGGCTTTT – 3′
bottom: 5′ – CGAAAAAAGCCAGGGAGTTTACACTTAGCTATGAATATTAACTAAGTGTAAACTCCCTGGCC – 3′

shRNA-2:

top: 5′ – ACCGGGGGAGTTTACACTTACGAAGTTAATATTCATAGCTTCGTAAGTGTAAACTCCCTTTT – 3′
bottom: 5′ – CGAAAAAAGGGAGTTTACACTTACGAAGCTATGAATATTAACTTCGTAAGTGTAAACTCCCC –3′

shRNA oligonucleotides were cloned into the DECIPHER pRSI12-U6-(sh)-HTS4-UbiC-TagRFP-2A-Puro lentiviral expression vector (Cellecta, cat. no. DVLIB-PS) following manufacturer’s instructions and then transformed into OneShot Top10 *E.coli* (Thermo Fisher, cat. no. C404003). Positives clones were verified by sequencing. 1 μg of two different shRNAs were packaged with lentiviral plasmids (MD2G, psPAX2) and transfected into HEK cells using JetPrime reagent (PolyPlus) according to manufacturer’s instructions. Media was collected 48 h after transfection and concentrated using Lenti-XTM concentrator (Takara Bio Europe SAS, cat. no. 631231) according to manufacturer’s instructions. Viral pellets were resuspended in 300 μl of DMEM and stored at −80°C.

SHSY5Y cells were seeded at 20% confluency in a 12 well plate before adding 10 μL lentivirus with shRNA. Cells were cultivated for 7 days with a media change on every third day. Cells were harvested by scraping, centrifuged for 5 min at 2000x g at 4°C, washed once with ice-cold 1X PBS and re-centrifuged for 5 min at 2000x g at 4°C. Samples were stored at −80°C until further use.

The knock down efficiency was analyzed using qPCR: RNA was extracted from the cell pellets using RNeasy Plus Mini Kit (QIAGEN, cat. no. 74134) according to the manufacturer’s protocol. A reverse transcription reaction was carried out to synthesize 250 ng cDNA using the High-Capacity cDNA Reverse Transcription Kit (Fisher Scientific, cat. no. 10400745), and qPCR was performed with the GoTaq® qPCR Master Mix (Promega, cat. no. A6001) following the manufacturer’s protocol on a Quant Studio 3 real-time PCR machine (Thermo Fisher).

### Subcellular Fractionation of mouse brain and cultured neurons

For subcellular fractionation, iPS-derived neurons were washed off the culture plate in ice-cold PBS, pelleted and fractionated using the Pierce Subcellular fractionation kit for cultured cells (Thermo Fisher) according to manufacturer’s instructions. Mouse brain samples were fractionated using the Pierce Subcellular fractionation kit for tissues (Thermo Fisher) according to manufacturer’s instructions. For analysis, 2-10 μl of each fraction were run on an SDS-PAGE and subjected to Western blotting as described above.

### Data analysis

Acquired LC-MS/MS data were processed using IsobarQuant (Franken, Mathieson et al., 2015) and Mascot (v2.2.07) with a reversed Uniprot homo sapiens database (UP000005640) including common contaminants. The following modifications were taken into account: Carbamidomethyl (C) (fixed modification), as well as Methyl (K), Dimethyl (K), Acetyl (K), Acetyl (N-term), Phospho (ST), Phospho (Y) and Oxidation (M) (variable modifications). The mass error tolerance for full scan MS spectra was set to 10 ppm and for MS/MS spectra to 0.5 Da. A maximum of 2 missed cleavages were allowed. A minimum of 2 unique peptides with a peptide length of at least seven amino acids and a false discovery rate below 0.01 were required on the peptide and protein level. Peptides with a MASCOT Score ≥ 32 were determined as reliable detected peptides (Morris, Knudsen et al., 2015).

Immunoblotting data were analyzed using Image Studio Lite (Li-cor Biosciences) and statistical analysis was performed with GraphPad Prism 7 (GraphPad Software) using the test noted within the respective figure legend.

## Acknowledgements

The study was funded by a contract research agreement between AbbVie Inc. (North Chicago, IL, USA) and BioMed X GmbH. Human brain samples were collected with informed consent by the London Neurodegenerative Diseases Brain Bank and the Southwest Dementia Brain Bank, members of the Brains for Dementia Research Network, and were provided in a strictly anonymized fashion. hiPSCs were derived from fibroblasts that are part of the NIA Aging Cell Repository at the Coriell Institute for Medical Research (Schondorf, Elschami et al., 2018). Informed consent was obtained by Coriell and the fibroblasts were provided in a strictly anonymized fashion for iPS derivation.

The authors thank Dr. Martin Fuhrmann for fruitful discussions and advice and Dr. Theron Johnson for access to the Meso Scale Discovery Quickplex platform.

## Author contributions

MB performed most experiments, coordinated co-author contributions and contributed to manuscript writing. NPO and EEH helped with SETD7 inhibitor and shRNA studies, BGR and VS performed antibody validation, DCS performed iPSC differentiation and neuronal subcellular fractionations. PH performed mass spectrometry and analyzed the resulting data. LG contributed brain tissue from tg4510 mice and manuscript writing. DEE supervised the project, performed fractionation experiments from mouse brain and wrote the manuscript. All authors contributed to data discussion and interpretation, read and approved the manuscript.

