## Supplementary material for "SETD7-mediated lysine monomethylation is abundant on non-hyperphosphorylated nuclear Tau": Suppl Fig legends

**Supplementary Figure S1:** **Methyl-Tau antibodies specifically recognize methylated over non-modified Tau peptides.** Non-modified recombinant Tau 2N4R, Tau 2N4R subjected to reductive methylation as well as methylated and non-methylated Tau-derived peptides were spotted onto nitrocellulose membranes and probed with the different methyl-Tau antibodies used in this study.

**Supplementary Figure S2: Three of five methyl-Tau antibodies demonstrate specificity for their methylation site in a cell lysate context.** HEK293T cells were transiently transfected with wt 2N4R Tau or mutant Tau proteins (K130R, K132R, K343R, K353R or K438R). While the antibodies directed against meK130, meK132 and meK353 specifically recognize wt, but not the corresponding mutant Tau, antibodies against meK343 and meK438 show no such specificity.

**Supplementary Figure S3:** **Treatment with λ-phosphatase abolishes the HMW Tau band.** Cytosolic and soluble nuclear fractions derived from 8- and 30-week old tg4510 mouse cortex were subjected to dephosphorylation with λ-phosphatase. Staining with Tau 12 demonstrates that the high molecular weight (HMW) Tau band apparent in 30-week old animals is abolished upon phosphatase treatment. Furthermore, the pT231 antibody staining, which strongly stains the HMW bands in untreated samples, is also removed by the treatment.

**Supplementary Figure S4: The influence of SETD7 inhibition or knockdown on total Tau levels. (A)** The treatment of SH-SY5Y cells with 5 µM (R)-PFI-2 for 24h does not alter total Tau levels as demonstrated by a representative Western blot and quantification. **(B)** Total Tau levels determined by electrochemiluminescence ELISA are also not altered in SH-SY5Y cells treated with 5 µM (R)-PFI-2 for 24h. **(C)** The treatment of iPS-derived neurons with 5 µM (R)-PFI-2 for 48h does not alter total Tau levels as demonstrated by a representative Western blot and quantification. **(D)** Total Tau levels determined by electrochemiluminescence ELISA are also not altered in iPS-derived neurons treated with 5 µM (R)-PFI-2 for 48h. **(E)** shRNA treatment against SETD7 leads to a slight reduction in total Tau levels in SH-SY5Y cells. For a representative Western blot, see Fig. 4C. **(F)** Total Tau levels do not significantly differ in HEK293T cells transiently transfected with wt 2N4R Tau or the K130R and K132R mutant proteins. For a representative Western blot, see Fig. 4D. Statistical significance was determined by Mann-Whitney test in **(A)**, **(B)**, **(C)**, **(D)** and **(E)**. For **(A)** – **(D)**, p>0.05, **: p<0.01. Statistical significance in **(F)** was determined by one-way ANOVA (ANOVA p=0.8096).
