## Supplementary material for "SETD7-mediated lysine monomethylation is abundant on non-hyperphosphorylated nuclear Tau": Suppl Figures

| | |
|---|---|
| 2N4R | recombinant tau |
| me2N4R | in vitro methylated recombinant tau |
| meK130 | CQARMVSK(me)SKDGTG-NH2 |
| meK132 | CRMVSKSK(me)DGTGSD-NH2 |
| meK343 | CVEVKSEK(me)LDFKDR-NH2 |
| meK353 | CKDRVQSK(me)IGSLDN-NH2 |
| meK438 | CDEVSASLAK(me)QGL-NH2 |
| K130/K132 | CQARMVSKSKDGTGSD-NH2 |
| K343 | CVEVKSEKLDFKDR-NH2 |
| K353 | CKDRVQSKIGDLDN-NH2 |
| K438 | CDEVSASLAKQGL-NH2 |

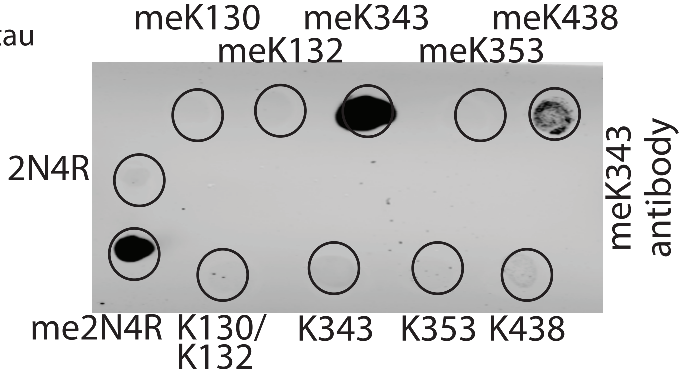


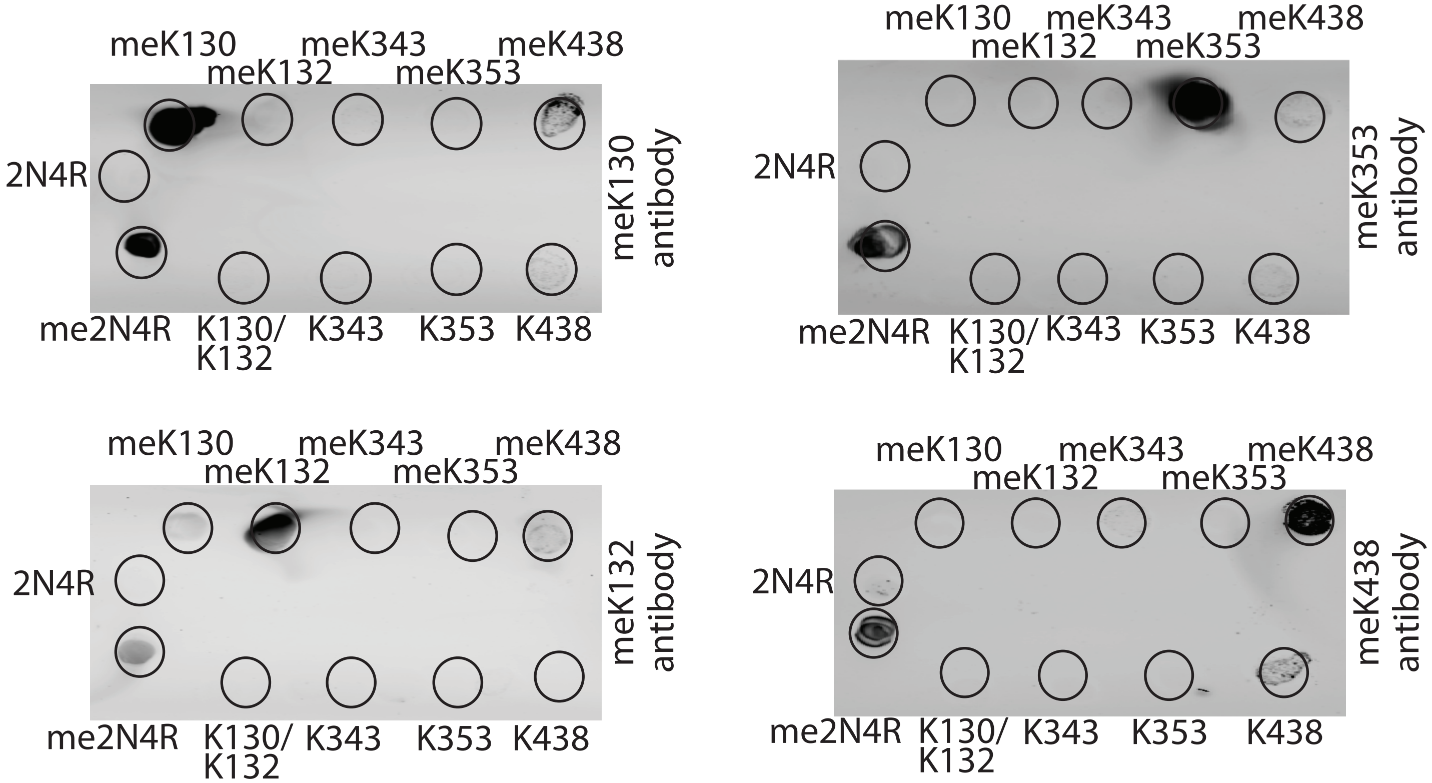


Suppl Figure S1

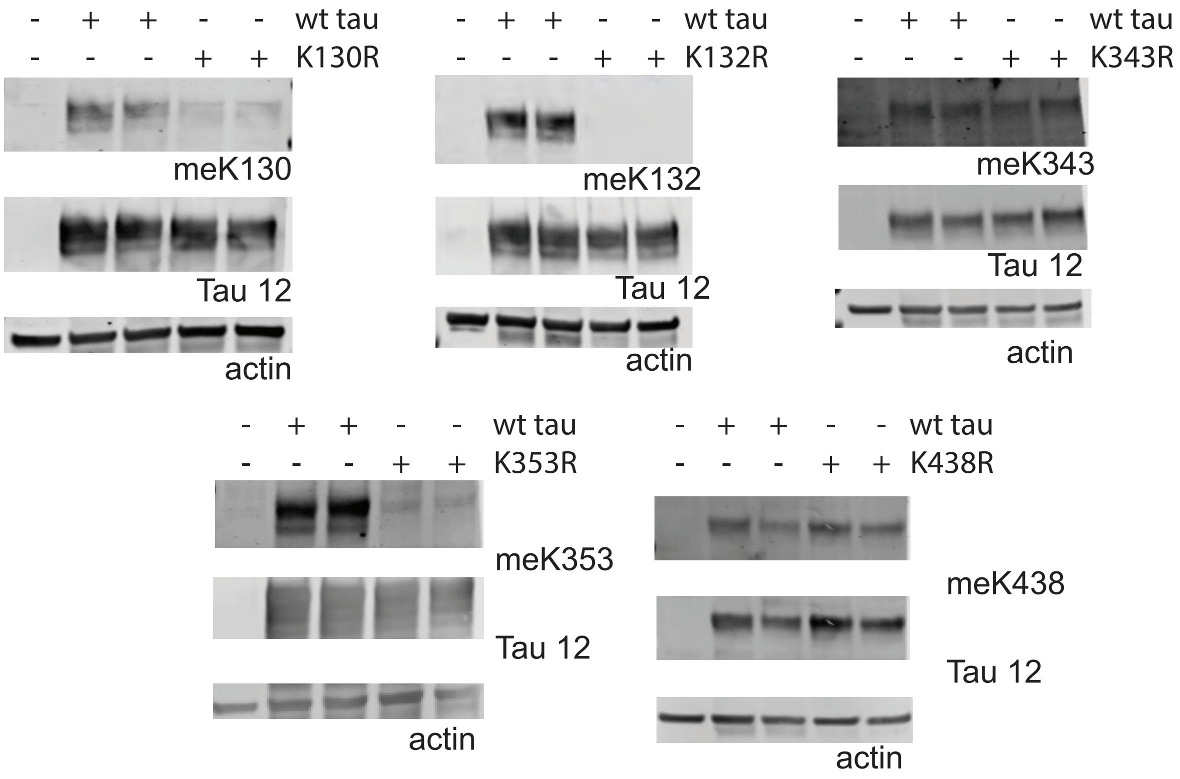


Suppl Figure S2

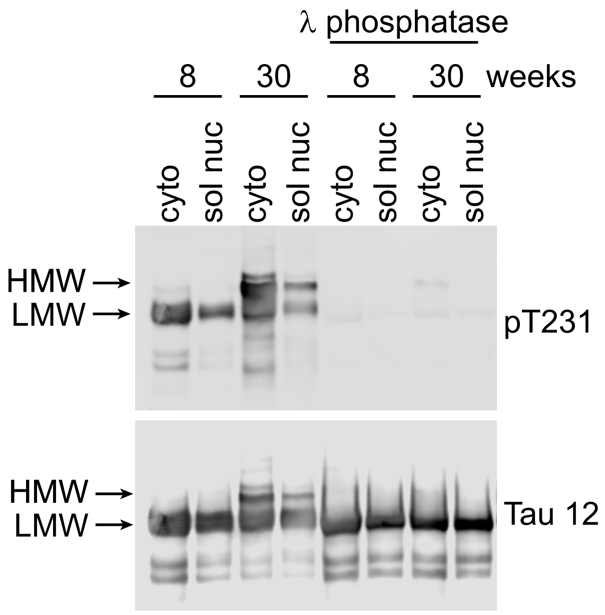


Suppl Figure S3

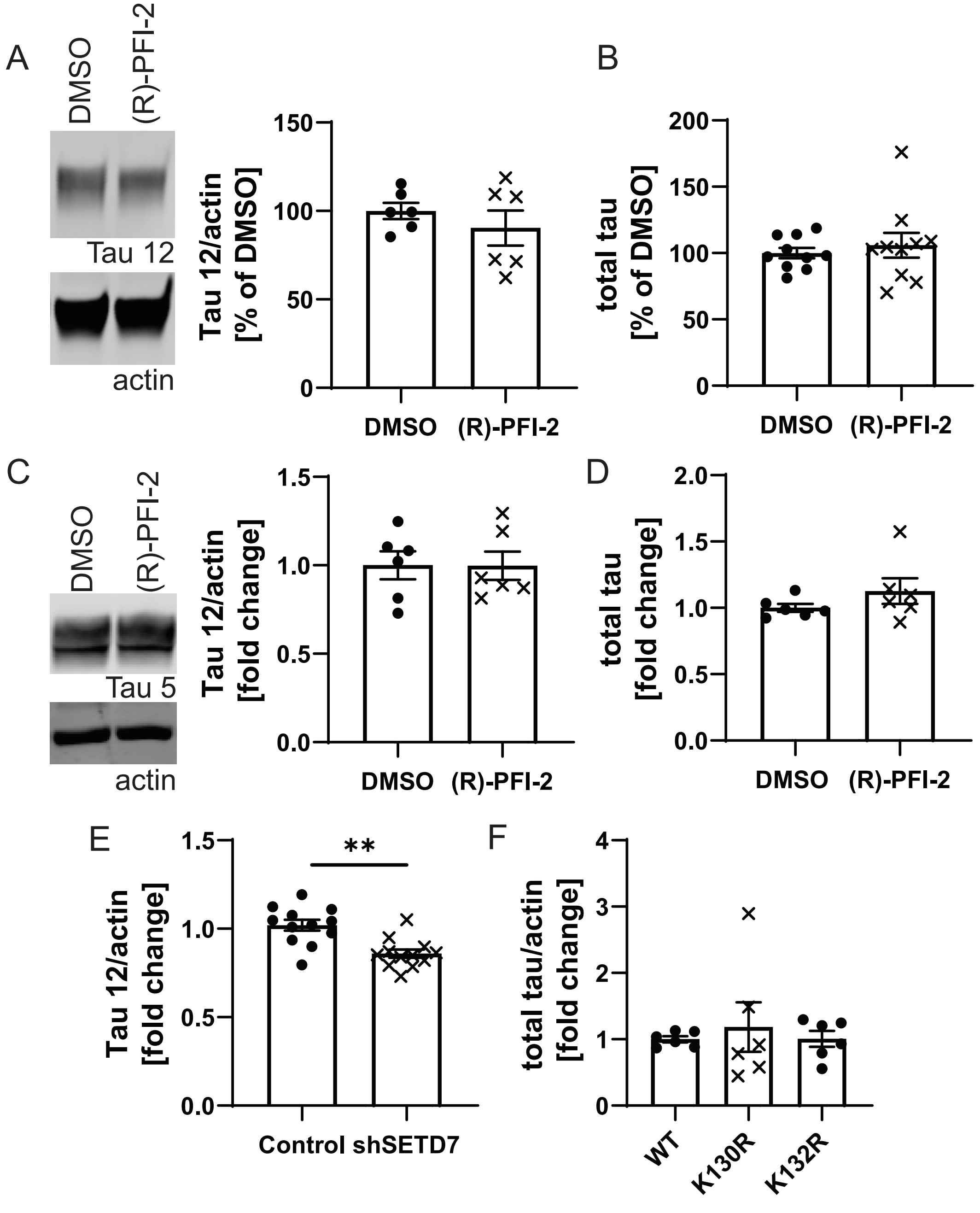


Suppl Figure S4
